# Optimizing Protein Tokenization: Reduced Amino Acid Alphabets for Efficient and Accurate Protein Language Models

**DOI:** 10.64898/2026.02.08.701987

**Authors:** Ella Rannon, David Burstein

**Affiliations:** The Shmunis School of Biomedicine and Cancer Research, Tel Aviv University

## Abstract

Protein language models (pLMs) typically tokenize sequences at the single-amino-acid level using a 20-residue alphabet, resulting in long input sequences and high computational cost. Sub-word tokenization methods such as Byte Pair Encoding (BPE) can reduce sequence length but are limited by the sparsity of long patterns in proteins encoded by the standard amino acid alphabet. Reduced amino acid alphabets, which group residues by physicochemical properties, offer a potential solution but their performances with sub-word tokenization have not been systematically studied. In this work, we investigate the combined use of reduced amino acid alphabets and BPE tokenization in protein language models. We pre-train RoBERTa-based pLMs de novo using multiple reduced alphabets and evaluate them across diverse downstream tasks. Our results show that reduced alphabets enable substantially shorter input sequences and faster training and inference. These findings suggest that alphabet reduction may facilitate more effective sub-word tokenization, enabling increased efficiency with marginal impact on predictive performance, and for specific tasks even improving accuracy.

## 1 Introduction

Advances in high-throughput sequencing have generated vast amounts of genomic sequence data, much of which remains unannotated (Sayers *et al*. 2025). The ability of language models (LMs) to learn complex patterns from large unlabeled corpora makes them well-suited to address this challenge (Bepler and Berger 2021; Iuchi *et al*. 2021; Ofer, Brandes and Linial 2021; Zhang *et al*. 2023; Oikonomou *et al*. 2024; Rannon and Burstein 2025). In particular, protein language models (pLMs) have demonstrated strong performance in capturing evolutionary, structural, and functional signals directly from amino acid sequences (Alley *et al*. 2019; Rao *et al*. 2019; Rives *et al*. 2021; Elnaggar *et al*. 2022; Lin *et al*. 2023).

A key design choice in pLMs is how protein sequences are segmented to basic language units called tokens (Testagrose and Boucher 2025). Many pLMs adopt character-level tokenization, treating individual amino acids as tokens (Alley *et al*. 2019; Heinzinger *et al*. 2019; Rao *et al*. 2019; Cao and Shen 2021; Rives *et al*. 2021; Lin *et al*. 2023). While this approach enables position-specific predictions and maintains a small vocabulary, it results in tokens encoding limited contextual information and long input sequences, leading to increased computational cost. Fixed-length *k*-mer tokenization (Asgari and Mofrad 2015; Kimothi *et al*. 2016) partially addresses this limitation but introduces sensitivity to insertions and deletions, severe vocabulary sparsity when utilizing a large k, and is based on the unrealistic assumption that informative sequence elements share a fixed length.

Sub-word tokenization methods, such as Byte Pair Encoding (BPE) (Sennrich, Haddow and Birch 2016), provide a flexible alternative by learning variable-length tokens from frequently co-occurring amino acid patterns. By treating protein sequences as raw symbol streams without predefined word boundaries, these methods are well-suited to biological sequences lacking linguistic structure. The resulting variable-length tokens can capture higher-order sequence features, including recurrent motifs of biological relevance, while reducing vocabulary sparsity and naturally handling unseen tokens. Sub-word tokenization has been successfully applied in several pLMs (Asgari, McHardy and Mofrad 2019; Wang *et al*. 2019; Elnaggar *et al*. 2022; Ferruz, Schmidt and Höcker 2022) and evaluated across vocabulary sizes (Suyunu, Taylan and Özgür 2024) and downstream tasks (Dotan *et al*. 2024; Tan *et al*. 2024). Nevertheless, its effectiveness is constrained by the sparsity of long patterns in the standard 20-amino-acid alphabet.

Protein sequences can be simplified by using reduced alphabets, where each character represents a group of amino acids with shared physico-chemical, functional, or structural properties (Liang et al. 2022). Reducing the amino acid alphabet can increase the frequency of longer recurring patterns with biological significance, enabling BPE to construct longer and more informative tokens. While reduced amino acid alphabets have been previously used in bioinformatics (Murphy, Wallqvist and Levy 2000; Weathers *et al*. 2004; Peterson *et al*. 2009; Ofer and Linial 2015), prior evaluations of these alphabets in pLMs focused exclusively on single-residue tokenization and reported reduced model performance (Ie-remie, Ewing and Niranjan 2024). The trade-off between alphabet resolution and token length has not yet been systematically explored on biological data. Combining reduced alphabets with sub-word tokenization may enable the construction of longer tokens while preserving or even enhancing the model’s ability to capture relevant biological patterns.

Here, we investigated the combined use of reduced amino acid alphabets and BPE sub-word tokenization in protein language models. We pretrained pLMs from scratch using different sizes of reduced alphabets and evaluated their performance across diverse downstream tasks. These include classification tasks for solubility, enzymes, transporter identification, two-component systems, protein-protein interactions (PPIs), as well as regression tasks for fluorescence, optimal temperature, and stability regression. We showed that reduced alphabets improved model performance on some tasks, while substantially reducing input sequence length and runtime. Our results highlight that incorporating prior knowledge of amino acid properties through alphabet reduction is an effective strategy for improving both the efficiency and, in certain tasks, the performance of protein language models.

## 2 Methods

### 2.1 Datasets

#### Corpus

This study’s corpus comprised protein sequences from assembled metagenomic contigs across various ecosystems from EBI’s Mgnify (Mitchell *et al*. 2020), together with proteins from assembled genomes and metagenomes in the NCBI GenBank Whole Genome Sequencing (WGS) database (Sayers *et al*. 2019), after filtering out Fungi, Metazoan, and Viridiplantae (downloaded on March 14, 2020). CD-HIT (Fu *et al*. 2012) (v4.6; -g 1 -s 0.8 -c 0.8) was utilized to reduce redundancy, retaining only the representative sequences from each cluster. The corpus was then split into test and training sets as follows. Genomes of isolated bacteria were assigned either to the training or test set, enforcing an 80:20 protein ratio. This ensured that proteins from the same genome do not appear in both the training and test sets. For metagenomes, samples from each ecosystem type were randomly split to reach an 80:20 protein ratio between the training and test sets within each ecosystem type.

#### Tokenizer training

For each alphabet, a BPE tokenizer was trained on a set of 28.3 million proteins from the corpus training set. Specifically, it included approximately 100,000 proteins randomly sampled from each ecosystem type in the corpus training set, as well as proteins from Ref-Seq’s representative and reference genomes, resulting in a total of 28,300,268 proteins. Vocabulary properties were assessed for each tokenizer on an evaluation set comprising ∼25,000 proteins per ecosystem type from the corpus test set, yielding 8,650,002 proteins.

#### Pre-training

Using the trained BPE tokenizer, a RoBERTa-based protein language model was pre-trained. The training set included a random subset of 15 million proteins from the corpus training set.

#### Homology detection

For this task, all protein sequences were obtained from UniRef90 (Suzek *et al*. 2015) and family annotations from InterPro (Paysan-Lafosse *et al*. 2023) (downloaded on August 3, 2025). The smallest 10% of families were removed. For each remaining family, 15 protein pairs were randomly sampled as positive examples. An approximately equal number of negative pairs was generated by sampling proteins from different families, ensuring each family appeared in 30 total pairs. The final dataset comprised 663,900 protein pairs.

To minimize potential data leakage between training and test sets, any test protein sharing more than 50% sequence identity at 80% or greater coverage with any training protein was removed using Linclust (Steinegger and Söding 2018) (--min-seq-id 0.5 -c 0.8; commit bad16c76).

#### Solubility and signal peptide prediction

The solubility dataset was acquired from DeepLocPro (Moreno *et al*. 2024) (v1.0), and contained 9,915 training and 1,991 test proteins after applying the 50% identity filtering to avoid potential leakage, as described above. The data was split into training and test in an 80:20 ratio using the original cross-validation partitions, with folds 0–3 taken as train, and fold 4 as test. The signal peptide training and test sets were obtained from SignalP (Teufel *et al*. 2022) (v6.0), resulting in 20,290 training and 2,021 test proteins after the same 50% identity filtering. In both cases, only a binary label was used.

#### Enzyme, transporter, and two-component system prediction

Kyoto Encyclopedia of Genes and Genomes (KEGG) (Kanehisa and Goto 2000) orthology (KO) identifiers were assigned to our corpus using the mapping from Miller, Stern, and Burstein (2022). Briefly, proteins associated with the same KO were subclustered using MMseqs2 cluster -s 7.5 -c 0.5 (Steinegger and Söding 2017), and each cluster was aligned with MAFFT (Katoh *et al*. 2002) and used to construct an HMM profile (Eddy 2011) for corpus annotation (Miller, Stern and Burstein 2022). Positive labels were assigned from the KEGG BRITE hierarchies representing enzymes, transporters, and two-component systems (ko01000, ko02000, and ko02022, respectively). Negative examples consisted of proteins annotated with KOs not belonging to the corresponding BRITE hierarchy. KOs were split into training, validation, and test sets in an 8:1:1 ratio based on their frequency in the data. Negative proteins were sampled to achieve a 10:1 negative-to-positive ratio. Due to the large resulting dataset size, stratified subsampling was applied, retaining 25% of the data for enzyme and transporter prediction tasks, and 30% for the two-component system prediction task. Finally, test proteins sharing ≥50% identity with any training protein were filtered out as described above. This process resulted in 4,235,473, 4,190,082, and 1,356,528 train proteins; 510,790, 497,336, and 208,524 validation proteins; and 505,100, 508,093, and 215,647 test proteins, for the enzyme, transporter, and two-component system prediction tasks, respectively.

#### PPI prediction

The dataset was based on B-PPI-DB (Agassy *et al*. 2025), including the *H. pylori* protein pairs used to test B-PPI (Agassy *et al*. 2025), which were processed as the other pairs in B-PPI-DB. The final PPI database included 162,703 training, 20,338 validation, and 20,338 test protein pairs. As this dataset construction process included Linclust clustering (--min-seq-id 0.4 -c 0.8), no further filtering was required.

#### Stability and Fluorescence prediction

The datasets were obtained from the TAPE benchmark (Rao *et al*. 2019) (downloaded on September 16, 2025). The stability and fluorescence datasets included 53,614 and 21,446 training proteins, 2,512 and 5,362 validation proteins, and 12,851 and 27,217 test proteins, respectively. The stability test set was reduced to 728 proteins after the 50% identity filtering. The fluorescence dataset was specifically included to assess model performance on closely related sequence variants. Since this dataset comprised variants of a single parent GFP sequence differing by only a few amino acids, the 50% identity filtering was not applied.

#### Optimal Temperature prediction

The dataset was downloaded from BRENDA (Hauenstein *et al*. 2025) (2025.1 release, version 1). Proteins with missing labels (“−999”) were removed. Sequences were clustered using MMseqs2 (Steinegger and Söding 2017) (-s 7.5 -c 0.8; commit bad16c76). Proteins labeled with a temperature range rather than a single value were removed, and the remaining proteins in their clusters were assigned to the test set. The sequences of all other clusters were assigned to the training set. This resulted in 6,991 train proteins and 2,591 test proteins after the 50% identity filtering of test proteins, as described above.

### 2.2 Tokenizer and Model Training

We evaluated five reduced amino acid alphabets of varying sizes based on different residue properties (Table 1). For each alphabet, a BPE tokenizer (Sennrich, Haddow and Birch 2016) was trained on the tokenizer dataset using a minimum token frequency of 2 and a vocabulary size of 5,000. This size was chosen to enable meaningful differences in token lengths across alphabets while remaining computationally tractable, as smaller vocabularies would yield uniformly short tokens and obscure the effects of alphabet reduction. The same vocabulary size was used across all alphabets to ensure a controlled comparison, allocating an equal “representational budget” and isolating the alphabet as the primary variable. A corresponding RoBERTa-based model (Liu *et al*. 2019) (termed Prot-BERTa_<alphabet size>) was then pre-trained, consisting of 12 attention heads, eight hidden layers, and a hidden dimension of 768. Each model was pre-trained on a single NVIDIA RTX A6000 GPU for five epochs with a batch size of 64, gradient accumulation of 8, and a maximum sequence length of 1,026 tokens, comprised of 1024 sequence tokens and two special tokens: <s> and </s> for marking the beginning and the end of the sequence, respectively. Pre-training was carried out using the masked language modeling (MLM) objective with a masking probability of 0.15.

**Table 1.** The amino acid alphabets used in this study and the groups of amino acids represented in their alphabet. The letter representing each group is written in parentheses.

| Alphabet Type | Size | Amino Acid Groups |
| --- | --- | --- |
| Amino Acids (Baseline) | 20 | A, C, D, E, F, G, H, I, K, L, M, N, P, Q, R, S, T, V, W, Y |
| Linclust’s Alphabet | 12 | AST (A), ND (N), EQ (E), FY (F), IV (I), KR (K), LM (L), C, G, H, P, W |
| Functional Groups | 8 | GVALI (S, Simple), ST (H, Hydroxy), CM (F, sulfur-containing), FY (M, aromatic), WHP (C, heterocyclic), NQ (N, amine-containing), DE (A, acidic), KR (B, basic) |
| Polarity | 4 | GAVLIFWMP (N, non-polar), STCYNQ (P, polar neutral), DE (S, negative charge), HKR (H, positive charge) |
| Hydrophilic-Hydrophobic | 2 | STNKEQHDRZB (L, hydrophilic), AGILMVPFWCY (B, hydrophobic) |

The pre-trained models were adapted on multiple downstream classification and regression tasks. For classification, a task-specific classification head was added, consisting of a dense layer with tanh activation applied to the final hidden state of the class token (<s>), followed by a linear projection layer. Models were trained using the AdamW optimizer, cross-entropy loss with a learning rate of 5⋅10^−5^ and a batch size 64, with gradient accumulation of 2. The models were trained for ten epochs, except for the PPI task, which was trained for two epochs.

For regression tasks, the output hidden states of the base model were mean-pooled and fed to a classification head, composed of a 256-dimensional linear layer with SiLU activation (Elfwing, Uchibe and Doya 2017), dropout with a rate of 0.15, and a projection layer. Models were trained using MAE loss with a learning rate of 2⋅10^−5^, weight decay of 0.01, warm-up ratio of 0.1, and a batch size of 64 for 15 epochs.

For all tasks except PPI prediction, the encoder weights were frozen to assess representation quality. In the PPI task, inputs were formatted as <s> protein1_tokens </s> </s> protein2_tokens </s>, which differs from the pre-training setup and therefore, all model parameters were fine-tuned.

### 2.3 Model Evaluation

The pre-trained embeddings from each ProtBERTa model were evaluated on the Diverse Genomic Embedding Benchmark (DGEB) (West-Roberts *et al*. 2024). Following the benchmark protocol, embeddings from the models’ middle (layer 4) and final (layer 7) layers were assessed, and the better-performing layer was reported for each task. The DGEB score was computed as the mean performance across tasks using their pre-defined primary metrics.

We further evaluated the quality of the models’ embedding using two tasks. For homology prediction, cosine similarity was calculated on mean-pooled embeddings for zero-shot classification of protein pairs as homologs. For signal peptide prediction, a kNN classifier (*k* = 5) based on cosine similarity between pooled embeddings was used, with classification scores defined by the proportion of positive neighbors.

Regression models were evaluated using mean squared error (MSE), mean absolute error (MAE), and root mean square error (RMSE), while binary classification models were evaluated using Area Under the ROC (AUROC), Area Under the Precision-Recall curve (AUPR), and maximal F1. For the two-component system prediction task, metrics were computed as macro averages across classes. Additionally, statistical significance between ProtBERTa_20 and each alternative model was assessed using McNemar’s test based on F1-optimizing predictions (or the class with maximal score for multiclass tasks).

Standard error was calculated for each metric using bootstrapping on the test set with *n* = 1,000 for all tasks, except for the homology, enzyme, transporter, and two-component system prediction tasks, where *n* = 100 was used instead due to the large size of the datasets.

### 2.4 Runtime Evaluation

Training and inference times were measured on a single NVIDIA RTX A6000 GPU. For each task, the test set was randomly subsampled ten times at multiple dataset sizes (increments of 1, 5, 10, 25, and 50), and inference runtime was recorded for the models’ inference on batches of 128 sequences. Prior to measurement, models were warmed up using 100 batches of size 128. For each dataset size, mean runtime and standard error were computed across the repeats. The dataset sizes of most tasks were scaled down by a factor of 1,000, except for the solubility/optimal temperature and stability prediction tasks, which were scaled down by factors of 39 and 500, respectively. Tokenizer training times and tokenization time were measured on a single process on an Intel Xeon Gold 6130 CPU. The tokenizer evaluation set was randomly subsampled ten times at multiple dataset sizes (5,000, 10,000, 25,000, 50,000, 75,000, and 100,000).

## 3 Results

### 3.1 Tokenizer Training

In this study, we investigated the merits and trade-offs of five different amino acid representations with different abstraction levels for protein language models (Table 1). Specifically, we explored:

1. The traditional 20-letter alphabet representing the 20 different amino acids.
2. A 12-letter representation based on the alphabet developed for the Linclust algorithm for fast clustering of proteins (Steinegger and Söding 2018).
3. An eight-letter representation based on the functional groups of the amino acids (Jain, Jain and Jain 2014).
4. A four-letter representation based on amino acids polarity (Ball, Hill and Scott 2014).
5. A two-letter representation where amino acids are split according to whether they are considered hydrophilic or hydrophobic.

We thus trained a BPE tokenizer (Sennrich, Haddow and Birch 2016) for each alphabet, resulting in different lengths of tokens (Table 2, Fig. 1A, Supplementary Fig. 1) and sentences (Fig. 1B, Supplementary Fig. 2, Supplementary Table 1). As the alphabet size decreases, recurring sequence patterns become more frequent, allowing BPE to merge characters into longer tokens. Thus, smaller alphabets yield more effective sequence compression and allow models to capture a broader context. This reduces running time and memory consumption, allowing utilization of more complex models (Vaswani *et al*. 2017; Korthikanti *et al*.). We pre-trained RoBERTa, a transformer-based language model (Liu *et al*. 2019), on a large corpus of microbial proteins from genomic and metagenomic samples using each tokenizer. The models were called ProtBERTa_X, where X denotes the relevant alphabet size (2, 4, 8, 12, or 20).

**Table 2.**
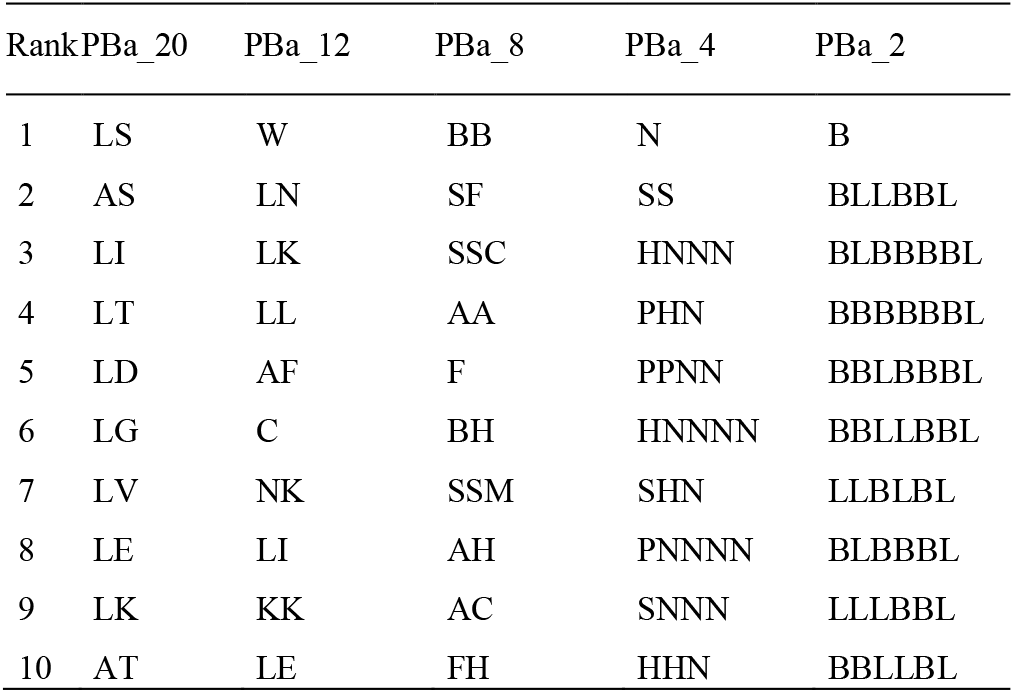
The ten most frequent tokens of each ProtBERTa (PBa) model observed in the tokenizer evaluation set.

**Fig. 1.**
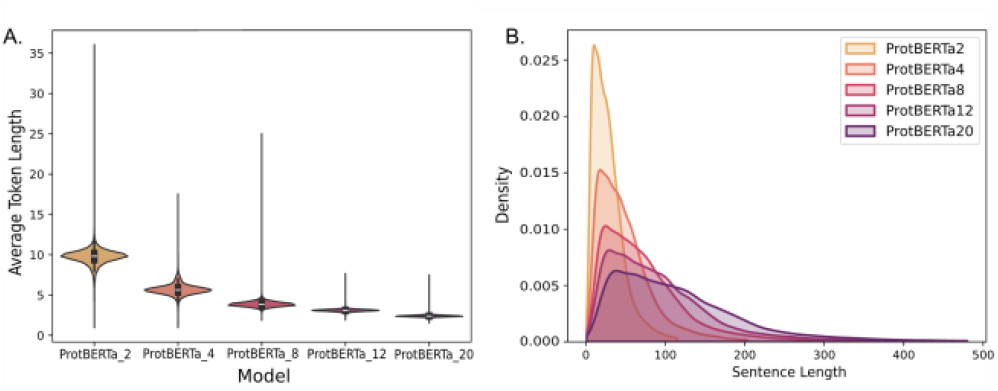
Properties of the BPE tokenizer trained on each amino acid alphabet. **A**. Average token length distribution for individual proteins. **B**. Distribution of tokenized sentence lengths (top 1% omitted for clarity).

**Fig. 2.**
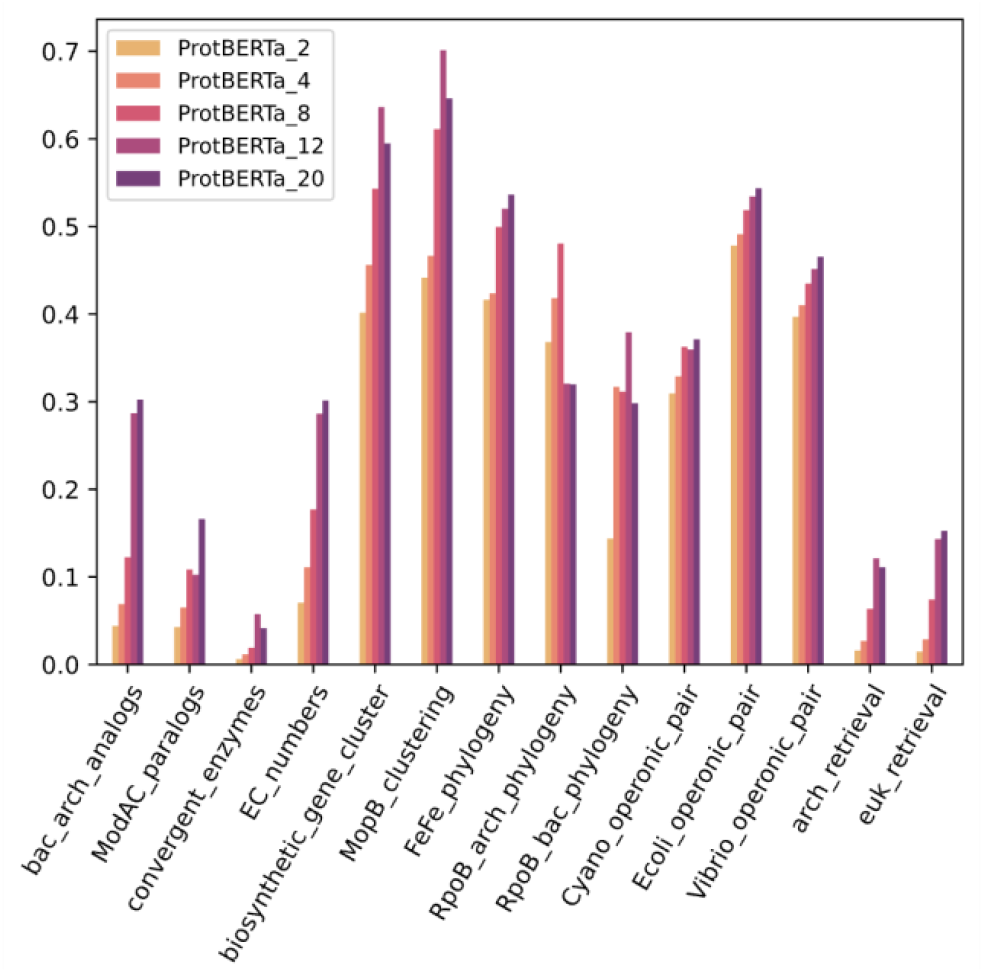
The performance of the different ProtBERTa models on primary metrics of the DGEB benchmark. The metrics descriptions are detailed in Supplementary Table 2.

### 3.2 Embedding Quality Assessment

#### 3.2.1 Evaluation on the Diverse Genomic Embedding Benchmark (DGEB)

To investigate the biological signals encoded in the embedding of each ProtBERTa model, we utilized the (DGEB) (West-Roberts *et al*. 2024), a predefined embedding benchmark for biological language models. DGEB includes six different tasks across 18 curated datasets, spanning sequences from all domains of life. The benchmark comprises both nucleic acid and amino acid modalities, but given the nature of our tokenizers, we have only utilized the amino acid benchmarking tasks (detailed in Supplementary Table 2). This benchmark is based on the model’s pre-trained embeddings directly.

The best overall DGEB benchmark score, which aggregates performance across all tasks, was achieved by ProtBERTa_12 (DGEB score of 0.35), followed closely by ProtBERTa_20 (0.347) and ProtBERTa_8 (0.309, Table 2). While ProtBERTa_12 achieved the highest aggregate score, ProtBERTa_20 was the best-performing model for the majority of individual tasks (eight out of 14, Fig. 2). In contrast, ProtBERTa_2 and ProtBERTa_4 were not the top models for any individual DGEB task.

#### 3.2.2 Zero-Shot Homology Prediction (pairwise binary classification)

We further evaluated the signals encoded in the models’ embeddings by zero-shot homology prediction, where the cosine similarity of the pretrained embeddings of pairs of proteins was used to train a model determining whether the two proteins belong to the same protein family. Even using raw embedding similarity alone, without any task-specific components, the classifiers based on the models’ embedding performed well (Fig. 3), with ProtBERTa_20 and ProtBERTa_12 achieving F1 scores of 0.818 and 0.813, respectively (Supplementary Table 3).

**Table 3.** DGEB scores of the different ProtBERTa (PBa) models. The best performance is marked in bold.

| Model | PBa_2 | PBa_4 | PBa_8 | PBa_12 | PBa_20 |
| --- | --- | --- | --- | --- | --- |
| DGEB Score | 0.225 | 0.259 | 0.309 | **0.35** | 0.347 |

**Fig. 3.**
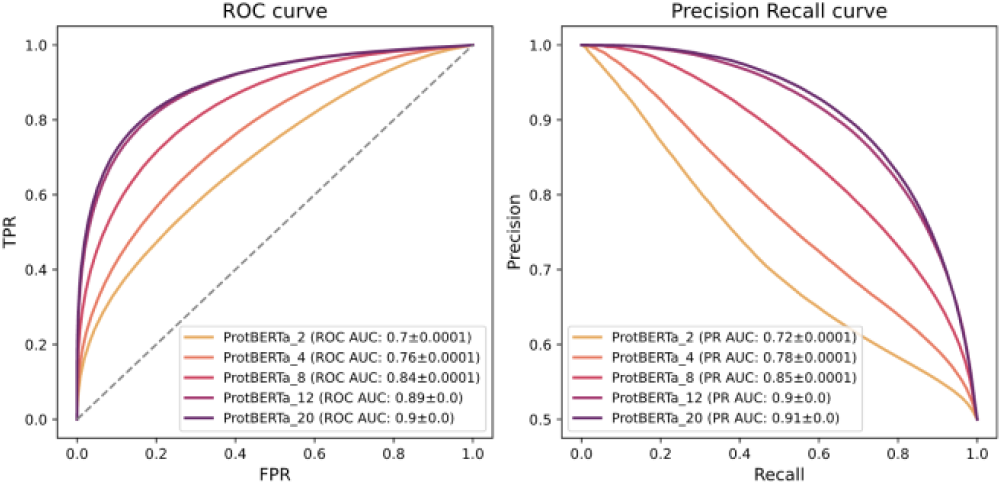
Zero-shot performance of the five ProtBERTa models on the task of homology detection. Left: ROC curves; right: precision–recall curves. The area under the curve score ± the standard error is noted in parentheses.

#### 3.2.3 Signal Peptide Detection (*k*-nearest neighbors)

We further assessed the models’ embeddings by training a *k*-nearest neighbors (kNN) classifier to identify signal peptides based on cosine similarity between the pre-trained embeddings. The labels of the test proteins were determined by majority vote among the *k* = 5 nearest train embeddings. ProtBERTa_20 achieved the best performance, with ProtBERTa_12 performing comparably (Fig. 4), while providing 1.28× compression (Supplementary Fig. 3).

**Fig. 4.**
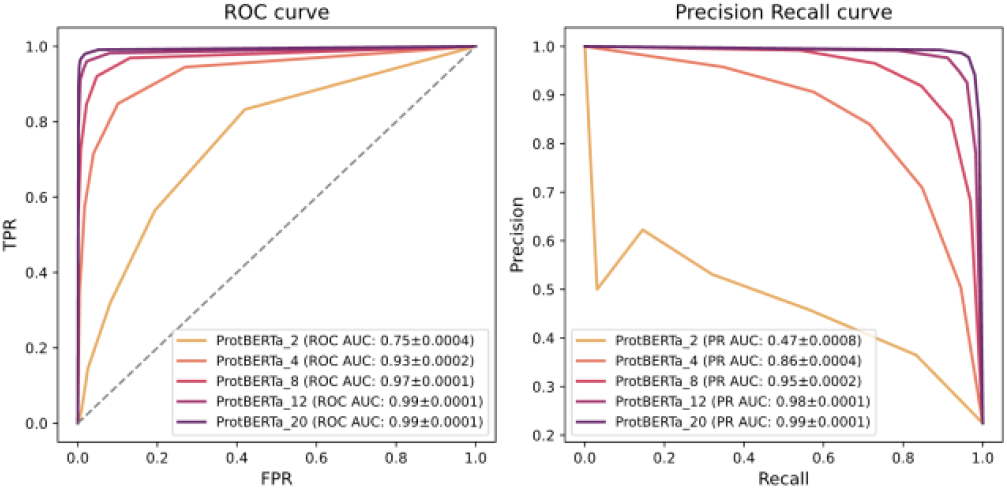
Performance of kNN classifiers using ProtBERTa pre-trained embeddings on the task of signal peptide classification. Left: ROC curves; right: precision–recall curves. The area under the curve score ± the standard error is written in parentheses.

### 3.3 Downstream Task Evaluation

#### 3.3.1 Classification Tasks

The ProtBERTa models were further adapted for a variety of downstream classification tasks. Specifically, task-specific classification heads were trained to predict protein solubility, identify enzymes, transporters, and two-component systems, and detect protein-protein interactions (PPIs). While most models performed well across the tasks, ProtBERTa_20 consistently exhibited the highest classification performance, while Prot-BERTa_12 and ProtBERTa_8 tended to achieve comparable performance (Fig. 5). In the solubility prediction task, where proteins were predicted as either soluble or membrane-bound, all models performed remarkably well, with ProtBERTa_8 achieving a slightly higher AUROC and AUPR (Supplementary Fig. 4, Supplementary Table 4), and ProtBERTa_20 marginally outperforming others in maximal F1. The performances of Prot-BERTa_12 and ProtBERTa_8 were not statistically different from the performance of ProtBERTa_20 (p-values of 0.057 and 0.175, respectively, McNemar’s test), while ProtBERTa_8 achieved over 1.5× input compression (Fig. 5).

**Table 4.** The relative task-specific training time of all the ProtBERTa models (PBa) compared to the baseline model, PBa_20. Values in parentheses indicate the relative input size (1/compression_ratio), which correlates with the runtime. TCS: Two-Component Systems.

| Task | PBa_12 | PBa_8 | PBa_4 | PBa_2 |
| --- | --- | --- | --- | --- |
| Solubility | 0.8 (0.77) | 0.69 (0.63) | 0.51 (0.43) | 0.33 (0.25) |
| Enzyme | 0.81 (0.78) | 0.67 (0.62) | 0.49 (0.43) | 0.32 (0.25) |
| Transporter | 0.78 (0.78) | 0.65 (0.62) | 0.47 (0.43) | 0.32 (0.25) |
| TCS | 0.79 (0.78) | 0.66 (0.62) | 0.47 (0.43) | 0.32 (0.25) |
| PPI | 0.79 (0.78) | 0.66 (0.63) | 0.46 (0.43) | 0.29 (0.25) |
| Stability | 0.96 (0.8) | 0.94 (0.69) | 0.89 (0.52) | 0.83 (0.35) |
| Temperature | 0.78 (0.78) | 0.64 (0.63) | 0.43 (0.43) | 0.25 (0.25) |
| Fluorescence | 0.86 (0.76) | 0.79 (0.62) | 0.63 (0.42) | 0.55 (0.24) |

**Fig. 5.**
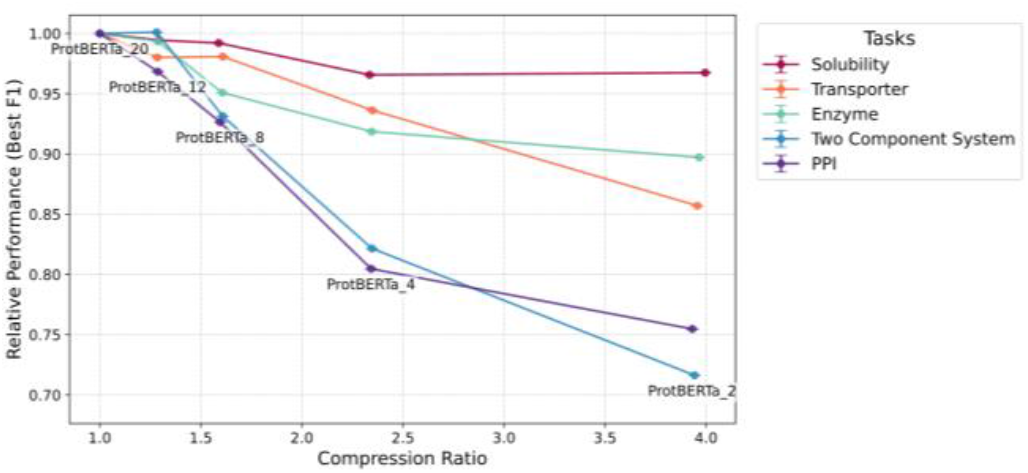
Performance vs. sentence length compression trade-off of the different Prot-BERTa models. The compression is calculated as average_sentence_length(Prot-BERTa_x)/average_sentence_length(ProtBERTa_20), while the relative performance is calculated as best_f1(ProtBERTa_x)/best_f1(ProtBERTa_20) for the binary classification models, and macro_f1(ProtBERTa_x)/macro_f1(ProtBERTa_20) for the two component classification task. The relative weighted F1 of the two-component system task is shown in Supplementary Figure 9.

For the enzyme detection task, ProtBERTa_20 and ProtBERTa_12 consistently achieved the strongest overall performance (Supplementary Figure 5, Supplementary Table 5). For the transporter task, ProtBERTa_20, ProtBERTa_12, and ProtBERTa_8 achieved comparable performance (Supplementary Figure 6, Supplementary Table 6). In both cases, Prot-BERTa_8 provided over 1.5× input compression with only modest performance degradation (2.5–5.5%, Fig. 5). In the two-component system multiclass task, where each protein is assigned as either a sensor, a response regulator, or neither, ProtBERTa_12 performed comparably to Prot-BERTa_20 (*p*-value of 0.354, McNemar’s test) with over 1.28× compression (Fig. 5, Supplementary Figure 7, Supplementary Table 7). In contrast, the protein–protein interaction task showed a clearer dependence on alphabet size, with performance decreasing as the alphabet was reduced (Supplementary Figure 8, Supplementary Table 8). Nevertheless, Prot-BERTa_8 still provided over 1.5× input compression with relatively limited performance loss (93% of ProtBERTa_20’s AUPR and best F1, Fig. 5).

#### 3.3.2 Regression Tasks

We also explored the performance of different ProtBERTa models on regression tasks, predicting protein stability, optimal temperature, and fluorescence levels. On the protein stability prediction task, the performance improved as the alphabet size increased, with ProtBERTa_4 demonstrating the best performance with an RMSE score of 0.543, and Prot-BERTa_20 following, with an RMSE score of 0.558 (Fig. 6A, Supplementary Table 9). Conversely, for the optimal temperature prediction task, model performance improves as the alphabet size decreases, with Prot-BERTa_2 outperforming the other models with an RMSE score of 17.577 (Fig. 6B, Supplementary Table 10). Finally, on the fluorescence prediction task, ProtBERTa_12 outperformed all the other models, achieving an RMSE score of 0.927 (Fig. 6C, Supplementary Table 11). While Prot-BERTa_20 and ProtBERTa_8 also performed relatively well.

**Fig. 6.**
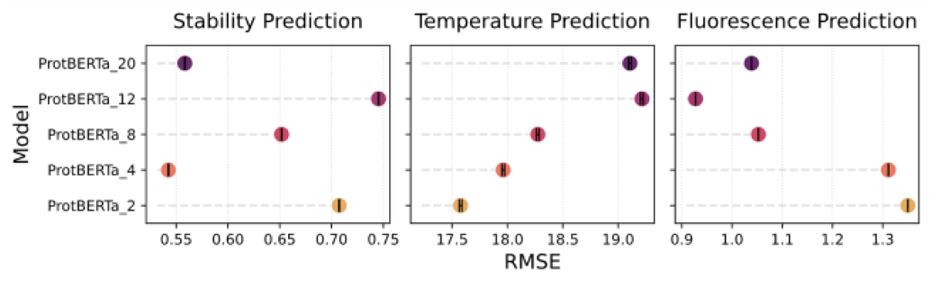
RMSE scores of the five ProtBERTa models on the regression tasks. **A**. Protein stability prediction. **B**. Optimal protein temperature prediction. **C**. Fluorescence prediction. Error bars represent standard error.

### 3.4 Runtime Comparison

Since the input sequence length directly affects computational cost, we hypothesized that the input compression achieved by reduced-alphabet tokenization would translate to shorter training and inference time. To verify this, we compared the running time of the models on downstream biological tasks described above. We observed that the decreases in training time roughly followed the models’ compression ratios. Specifically, the Prot-BERTa_4 required approximately half the training time of Prot-BERTa_20, while ProtBERTa_2 required roughly one third (Table 3). We observed a similar trend for the tokenizer training time (Supplementary Figure 10). In contrast, tokenization times were nearly identical across ProtBERTa_20, ProtBERTa_12, and ProtBERTa_8, with ProtBERTa_2 having slightly lower tokenization times (Supplementary Figure 11).

This trend is consistent with the computational structure of transformer models, which comprise self-attention layers with *O*(*s*^2^*d*) complexity and feed-forward layers with *O*(*sd*^2^) complexity, where *s* is the sequence length and *d* is the hidden dimension size. Since most tokenized sequences in our datasets are shorter than 200 tokens, with a mean length below 123 tokens (Fig. 1B, Supplementary Table 1), and the models’ hidden dimension size is 768, *d* has a greater contribution to the overall computation. Consequently, the feed-forward layers are expected to have a greater impact on runtime than the self-attention layers. This may explain the approximately linear scaling of runtime with input length, and the observed reductions in training time.

We note that, except for the PPI model, which was fully fine-tuned, all encoder parameters were frozen, with only the task-specific heads updated. In addition, the stability prediction dataset contains unusually short proteins (mean length of 43.2 amino acids), which likely accounts for the smaller-than-expected runtime differences observed for this task relative to others.

We also compared the inference time of the models according to the test set sample size. We observed that a similar trend also occurred during the inference stage, where the models exhibited approximately the same runtime ratio across different sizes of datasets, with ProtBERTa_20 running about four times longer than ProtBERTa_2. A linear relationship between the runtime and number of sequences was observed across the different tasks and models (Fig. 7).

**Fig. 7.**
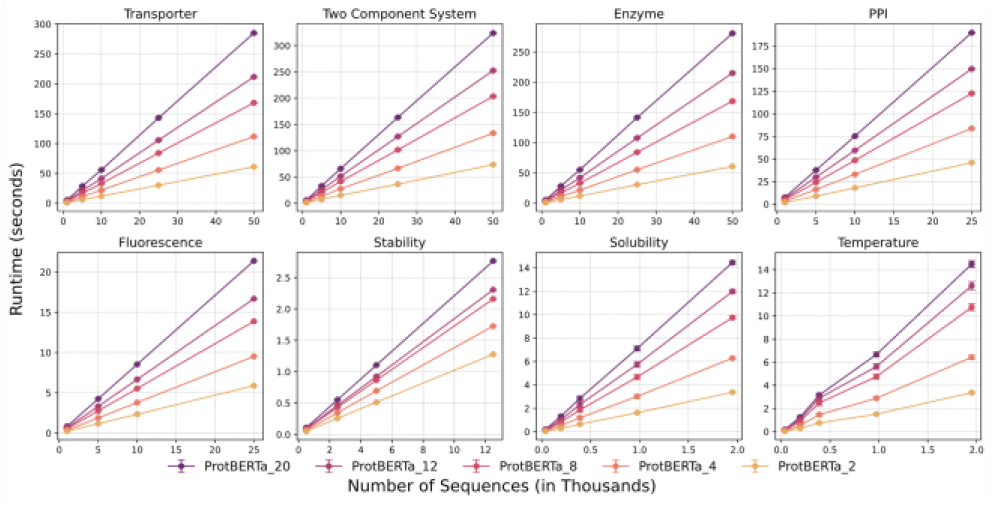
Inference run time comparison for the five ProtBERTa models on a variety of downstream tasks. Inference time is measured as a function of the dataset size (in thousands). Error bars represent standard error.

## 4 Conclusion

Although reduced amino acid alphabets have been widely used in bioinformatics (Murphy, Wallqvist and Levy 2000; Weathers *et al*. 2004; Peterson *et al*. 2009; Ofer and Linial 2015) and previously evaluated in the context of protein language models (Ieremie, Ewing and Niranjan 2024), they had not been combined with sub-word tokenization to improve pLM efficiency. Our results indicate that, while the full 20-amino-acid alphabet achieves the highest performance on most tasks, the models trained on the reduced alphabets have outperformed the baseline model on specific tasks. Notably, models trained on reduced alphabets typically incurred only minor performance loss, while providing substantial reductions in training and inference times.

We hypothesize that the decreased performance of models trained with reduced alphabets on classification tasks stems from the loss of fine-grained biochemical information encoded by specific amino acids. This effect is particularly pronounced in the protein–protein interaction (PPI) prediction task, where precise residue identities are critical for mediating physical interactions. In contrast, for optimal temperature prediction, performance improves as the alphabet size decreases. This may reflect the relatively small dataset combined with high label variability, making the task inherently challenging; under such conditions, learning more generalized representations encoding global thermodynamic signatures that drive thermal stability, while filtering out sequence-specific noise that can lead to overfitting, could be advantageous. The fluorescence and stability prediction tasks represent a middle ground, where an intermediate alphabet (12 characters for fluorescence, 4 for stability) performs best, possibly because it captures the necessary evolutionary constraints of the protein fold while maintaining enough relevant chemical details. Nevertheless, our findings do not allow for a definitive explanation of why certain alphabet sizes are better suited to specific tasks. We therefore recommend evaluating multiple alphabet configurations when applying subword tokenization-based protein language models, as reduced alphabets can out-perform standard representations in certain settings, while in others they achieve comparable performance with the added benefit of reduced computational cost.

We note that while a full-alphabet model could, in principle, learn similar generalizations from sufficient labeled training data, in practice, this may be difficult to achieve. A model pre-trained extensively on the full 20-amino acid alphabet develops rich, fine-grained representations that distinguish between all amino acids. When applied to a downstream task with a small dataset, such distinctions are noise rather than signal; the model must effectively unlearn these pre-trained representations, a process that requires substantial data and may not converge when limited labeled data are available. A reduced-alphabet model, by contrast, never encodes these distinctions in the first place, providing representations that are already at the appropriate resolution for the task. This is analogous to the choice of amino acids over codons as the basic unit of protein modeling.

To ensure a fair comparison, all models were pre-trained and applied to downstream tasks using identical hyperparameters. While task-specific and model-specific hyperparameter optimization could alter performance trends and potentially reduce differences between models, we avoided this approach to better isolate the effects of alphabet reduction and the associated physicochemical properties. Future work could explore model-specific optimization strategies as well as additional reduced alphabets.

It is worth noting that our study focuses on the impact of integrating reduced amino acid alphabets within subword tokenization for pLMs. Accordingly, our primary baseline is a 20-amino acid BPE tokenizer, rather than the character-level pLMs commonly used in prior work. Character-level models are typically more appropriate for residue-level biological tasks, for which BPE-based tokenization is less suitable. Reduced alphabets may also obscure mutations between amino acids within the same group, limiting applications such as variant effect prediction. These tasks would therefore benefit from alternative tokenization strategies.

A natural direction for future work is a systematic comparison between reduced-alphabet BPE models, character-level baselines, and existing pLMs in terms of performance and efficiency. Additionally, comparing our approach against alternative efficiency strategies, such as knowledge distillation or smaller architectures, would be valuable.

Furthermore, our experiments were conducted using a relatively small RoBERTa-based pLM. We expect similar trends to hold for larger models, particularly with respect to speed improvements in attention-based architectures, as the degree of sequence compression is determined by the tokenizer. However, further investigation is needed to assess the effects of reduced alphabets and BPE tokenization across models of varying architectures and scales.

Overall, our findings highlight the merits of combining reduced amino acid alphabets with sub-word tokenization as an effective strategy for compressing pLM input sequences, enabling more efficient training and inference while preserving biologically meaningful signals.

## Acknowledgements

We thank Asaf Zorea, Chen Agassy, Guy Shur, and Dr. Karin Mittelman for their feedback during this research.

## Funding

This work has been supported by the Israel Science Foundation [grant number 355/23], a grant from the Tel Aviv University Center for AI and Data Science (TAD), and by a fellowship to ER from the Edmond J. Safra Center for Bioinformatics at Tel Aviv University.

### Conflict of Interest

none declared.

## Data Availability

The datasets used in this study were obtained from publicly available sources, including EMBL-EBI MGnify (https://www.ebi.ac.uk/metagenomics) and NCBI Whole Genome Shotgun (WGS) (https://www.ncbi.nlm.nih.gov/genbank/wgs). The code supporting this work is publicly available on GitHub at https://github.com/burstein-lab/BioTokenizers. The trained tokenizers and pretrained models are available via Zenodo at https://doi.org/10.5281/zenodo.18256943 and https://doi.org/10.5281/zenodo.18257091.

## Notes

### Competing Interest Statement

The authors have declared no competing interest.

### Summary of Updates

The test datasets have been revised, and sequences with more than 50% identity to any of the train proteins were removed.

## References

Agassy C, Samuel B, Mayo S et al. B-PPI: A Cross-Attention Model for Large-Scale Bacterial Protein-Protein Interaction Prediction. 2025:2025.12.23.696145.

Alley EC, Khimulya G, Biswas S et al. Unified rational protein engineering with sequence-based deep representation learning. Nat Methods 2019;16:1315–22.

Asgari E, McHardy AC, Mofrad MRK. Probabilistic variable-length segmentation of protein sequences for discriminative motif discovery (DiMotif) and sequence embedding (ProtVecX). Sci Rep 2019;9:3577.

Asgari E, Mofrad MRK. Continuous Distributed Representation of Biological Sequences for Deep Proteomics and Genomics. PLOS ONE 2015;10:e0141287.

Ball DW, Hill JW, Scott RJ. 18.1: Properties of Amino Acids. The Basics of General, Organic, and Biological Chemistry. LibreTexts, 2014.

Bepler T, Berger B. Learning the protein language: Evolution, structure, and function. Cell Systems 2021;12:654–669.e3.

Cao Y, Shen Y. TALE: Transformer-based protein function Annotation with joint sequence–Label Embedding. Bioinformatics 2021;37:2825–33.

Dotan E, Jaschek G, Pupko T et al. Effect of tokenization on transformers for biological sequences. Bioinformatics 2024;40:btae196.

Eddy SR. Accelerated Profile HMM Searches. PLOS Computational Biology 2011;7:e1002195.

Elfwing S, Uchibe E, Doya K. Sigmoid-Weighted Linear Units for Neural Network Function Approximation in Reinforcement Learning. 2017, DOI: 10.48550/arXiv.1702.03118.

Elnaggar A, Heinzinger M, Dallago C et al. ProtTrans: Toward Understanding the Language of Life Through Self-Supervised Learning. IEEE Trans Pattern Anal Mach Intell 2022;44:7112–27.

Ferruz N, Schmidt S, Höcker B. ProtGPT2 is a deep unsupervised language model for protein design. Nat Commun 2022;13:4348.

Fu L, Niu B, Zhu Z et al. CD-HIT: accelerated for clustering the next-generation sequencing data. Bioinformatics 2012;28:3150–2.

Hauenstein J, Jeske L, Jäde A et al. BRENDA in 2026: a Global Core Biodata Resource for functional enzyme and metabolic data within the DSMZ Digital Diversity. Nucleic Acids Res 2025:gkaf1113.

Heinzinger M, Elnaggar A, Wang Y et al. Modeling aspects of the language of life through transfer-learning protein sequences. BMC Bioinformatics 2019;20:723.

Ieremie I, Ewing RM, Niranjan M. Protein language models meet reduced amino acid alphabets. Bioinformatics 2024;40:btae061.

Iuchi H, Matsutani T, Yamada K et al. Representation learning applications in biological sequence analysis. Computational and Structural Biotechnology Journal 2021;19:3198–208.

Jain JL, Jain S, Jain N. Proteins I - General Structure. Fundamentals of Biochemistry. 7th edn. S Chand and Company Ltd., 2014, 172.

Kanehisa M, Goto S. KEGG: Kyoto Encyclopedia of Genes and Genomes. Nucleic Acids Research 2000;28:27–30.

Katoh K, Misawa K, Kuma K et al. MAFFT: a novel method for rapid multiple sequence alignment based on fast Fourier transform. Nucleic Acids Res 2002;30:3059–66.

Kimothi D, Soni A, Biyani P et al. Distributed Representations for Biological Sequence Analysis. 2016.

Korthikanti V, Casper J, Lym S et al. REDUCING ACTIVATION RECOMPUTATION IN LARGE TRANSFORMER MODELS.

Liang Y, Yang S, Zheng L et al. Research progress of reduced amino acid alphabets in protein analysis and prediction. Computational and Structural Biotechnology Journal 2022;20:3503–10.

Lin Z, Akin H, Rao R et al. Evolutionary-scale prediction of atomic-level protein structure with a language model. Science 2023;379:1123–30.

Liu Y, Ott M, Goyal N et al. RoBERTa: A Robustly Optimized BERT Pretraining Approach. 2019, DOI: 10.48550/arXiv.1907.11692.

Miller D, Stern A, Burstein D. Deciphering microbial gene function using natural language processing. Nat Commun 2022;13:5731.

Mitchell AL, Almeida A, Beracochea M et al. MGnify: the microbiome analysis resource in 2020. Nucleic Acids Res 2020;48:D570–8.

Moreno J, Nielsen H, Winther O et al. Predicting the subcellular location of prokaryotic proteins with DeepLocPro. 2024:2024.01.04.574157.

Murphy LR, Wallqvist A, Levy RM. Simplified amino acid alphabets for protein fold recognition and implications for folding. Protein Engineering 2000;13:149–52.

Ofer D, Brandes N, Linial M. The language of proteins: NLP, machine learning & protein sequences. Computational and Structural Biotechnology Journal 2021;19:1750–8.

Ofer D, Linial M. ProFET: Feature engineering captures high-level protein functions. Bioinformatics 2015;31:3429–36.

Oikonomou ED, Karvelis P, Giannakeas N et al. How natural language processing derived techniques are used on biological data: a systematic review. Netw Model Anal Health Inform Bioinforma 2024;13:23.

Paysan-Lafosse T, Blum M, Chuguransky S et al. InterPro in 2022. Nucleic Acids Res 2023;51:D418–27.

Peterson EL, Kondev J, Theriot JA et al. Reduced amino acid alphabets exhibit an improved sensitivity and selectivity in fold assignment. Bioinformatics 2009;25:1356–62.

Rannon E, Burstein D. Leveraging Natural Language Processing to Unravel the Mystery of Life: A Review of NLP Approaches in Genomics, Transcriptomics, and Proteomics. 2025, DOI: 10.48550/arXiv.2506.02212.

Rao R, Bhattacharya N, Thomas N et al. Evaluating Protein Transfer Learning with TAPE. Advances in Neural Information Processing Systems. Vol 32. Curran Associates, Inc., 2019.

Rives A, Meier J, Sercu T et al. Biological structure and function emerge from scaling unsupervised learning to 250 million protein sequences. Proceedings of the National Academy of Sciences 2021;118:e2016239118.

Sayers EW, Cavanaugh M, Clark K et al. GenBank. Nucleic Acids Research 2019:gkz956.

Sayers EW, Cavanaugh M, Frisse L et al. GenBank 2025 update. Nucleic Acids Res 2025;53:D56–61.

Sennrich R, Haddow B, Birch A. Neural Machine Translation of Rare Words with Subword Units. In: Erk K, Smith NA (eds), Proceedings of the 54th Annual Meeting of the Association for Computational Linguistics (Volume 1: Long Papers). Berlin, Germany: Association for Computational Linguistics, 2016, 1715–25.

Steinegger M, Söding J. MMseqs2 enables sensitive protein sequence searching for the analysis of massive data sets. Nat Biotechnol 2017;35:1026–8.

Steinegger M, Söding J. Clustering huge protein sequence sets in linear time. Nat Commun 2018;9:2542.

Suyunu B, Taylan E, Özgür A. Linguistic Laws Meet Protein Sequences: A Comparative Analysis of Subword Tokenization Methods. 2024, DOI: 10.48550/arXiv.2411.17669.

Suzek BE, Wang Y, Huang H et al. UniRef clusters: a comprehensive and scalable alternative for improving sequence similarity searches. Bioinformatics 2015;31:926–32.

Tan Y, Li M, Zhou Z et al. PETA: evaluating the impact of protein transfer learning with sub-word tokenization on downstream applications. J Cheminform 2024;16:92.

Testagrose C, Boucher C. Tokenization and deep learning architectures in genomics: A comprehensive review. Computational and Structural Biotechnology Journal 2025;27:3547–55.

Teufel F, Almagro Armenteros JJ, Johansen AR et al. SignalP 6.0 predicts all five types of signal peptides using protein language models. Nat Biotechnol 2022;40:1023–5.

Vaswani A, Shazeer N, Parmar N et al. Attention Is All You Need. 2017, DOI: 10.48550/arXiv.1706.03762.

Wang Y, You Z-H, Yang S et al. A High Efficient Biological Language Model for Predicting Protein–Protein Interactions. Cells 2019;8:122.

Weathers EA, Paulaitis ME, Woolf TB et al. Reduced amino acid alphabet is sufficient to accurately recognize intrinsically disordered protein. FEBS Letters 2004;576:348–52.

West-Roberts J, Kravitz J, Jha N et al. Diverse Genomic Embedding Benchmark for functional evaluation across the tree of life. 2024:2024.07.10.602933.

Zhang S, Fan R, Liu Y et al. Applications of transformer-based language models in bioinformatics: a survey. Bioinformatics Advances 2023;3:vbad001.

