## Supplementary Materials for "Optimizing Protein Tokenization: Reduced Amino Acid Alphabets for Efficient and Accurate Protein Language Models"

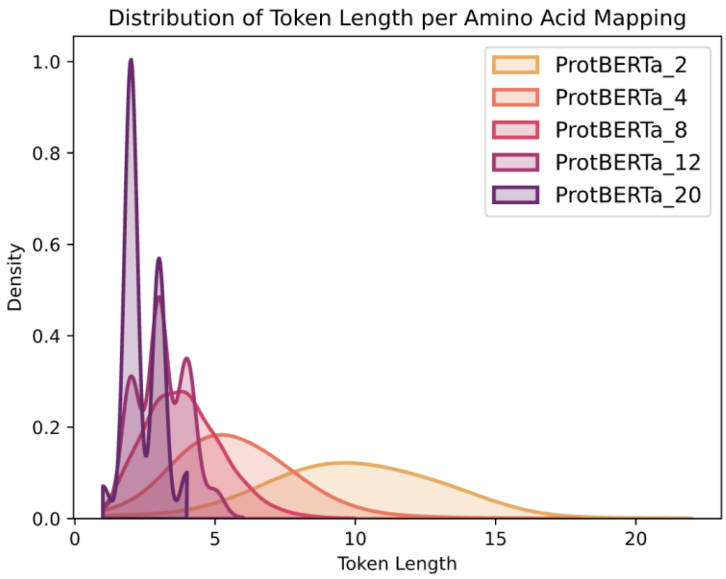


**Supplementary Figure 1. Token length distribution of the test tokenizer dataset for each reduced alphabet**. For clarity, only lengths up to the 0.999$^{th}$ quantile are presented.

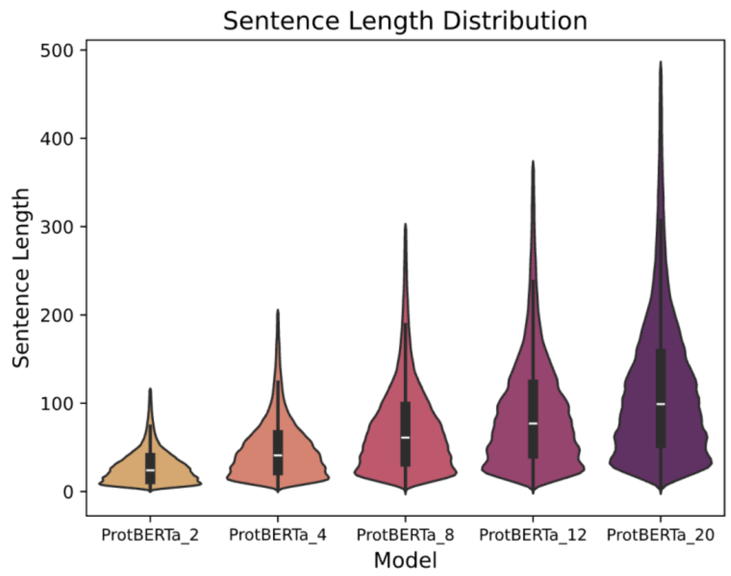


**Supplementary Figure 2. Sentence length distribution of the test tokenizer dataset for each reduced alphabet.** For clarity, only lengths up to the 0.99$^{th}$ quantile are presented.

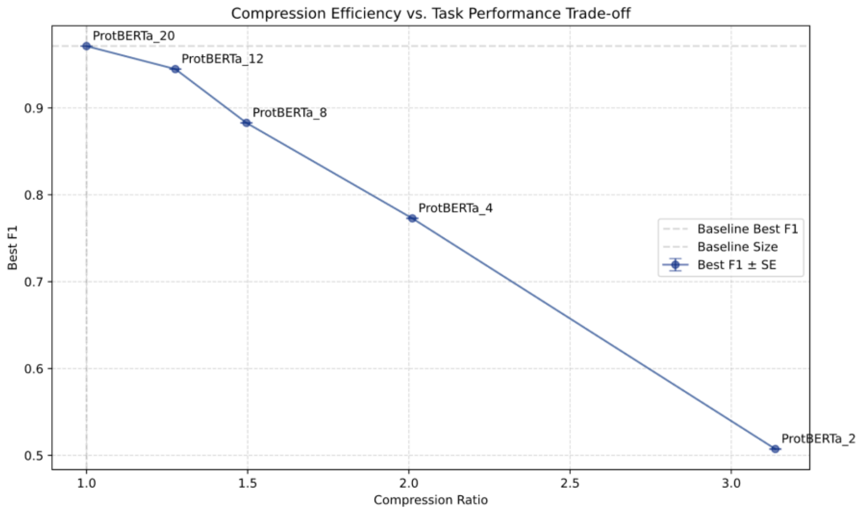


**Supplementary Figure 3. Trade-off between the performance of a kNN classifier trained on ProtBERTa's embeddings.** The performance is measured in terms of the best F1 and sentence length compression of the different ProtBERTa models on the task of signal peptide prediction. The compression is calculated as the average sentence length ratio between the alternative models and ProtBERTa_20. Error bars represent standard error.

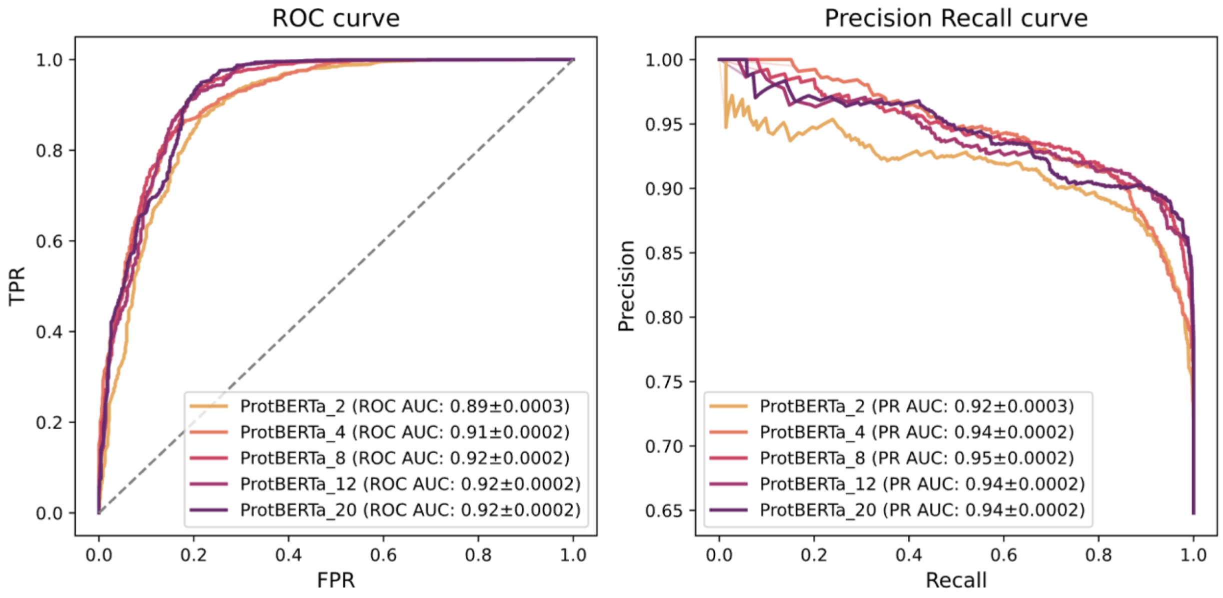


**Supplementary Figure 4**. **Performance of the five ProtBERTa models applied to the solubility prediction task.** Left: ROC curve, right: precision-recall curve. The area-under-the-curve (AUC) score and standard error for each model appear in parentheses.

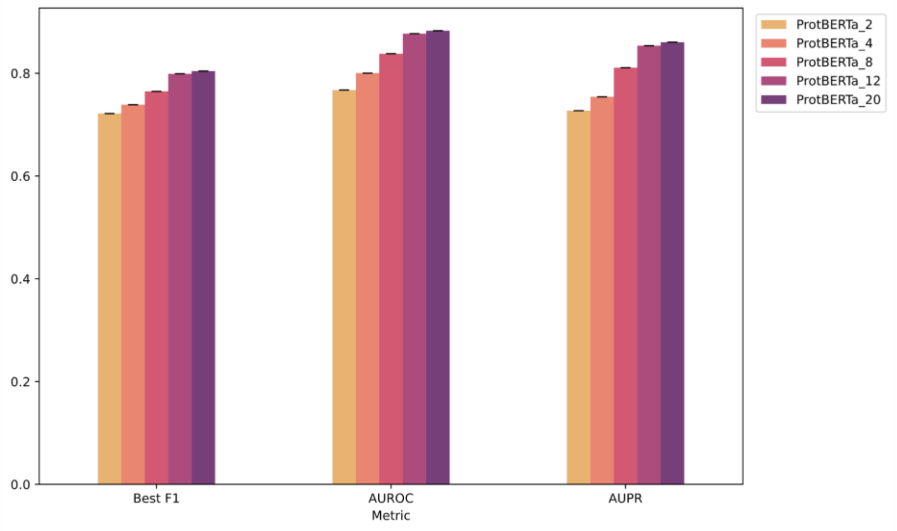


**Supplementary Figure 5. Performance of the five ProtBERTa models on the task of enzyme detection.** Standard error is shown as black error bars.

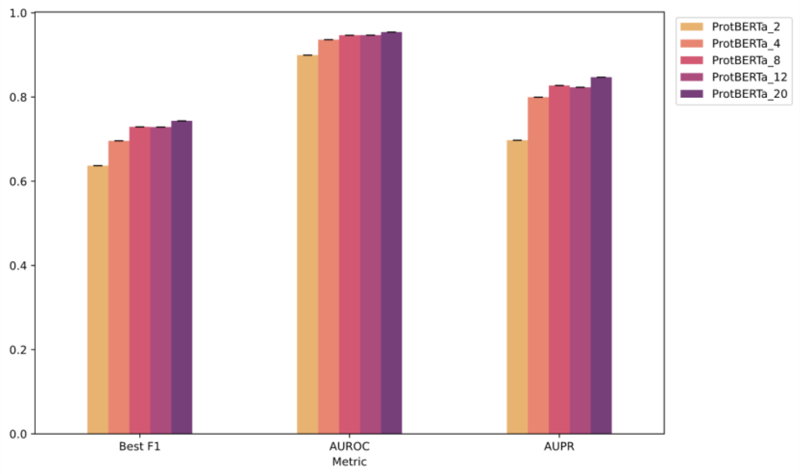


**Supplementary Figure 6. Performance of the five ProtBERTa models on the task of transporter detection.** Standard error is shown as black error bars.

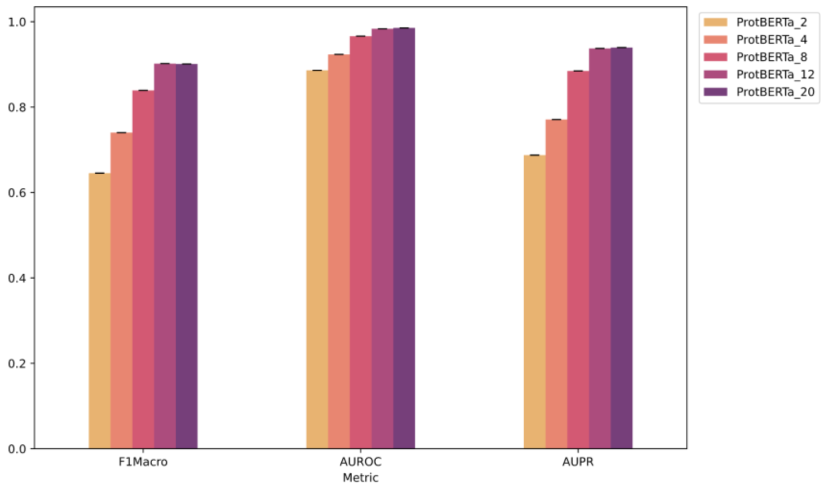


**Supplementary Figure 7. Performance of the five ProtBERTa models on the task of two-component system prediction.** All the metrics are calculated using a macro average across the three classes. Standard error is shown as black error bars.

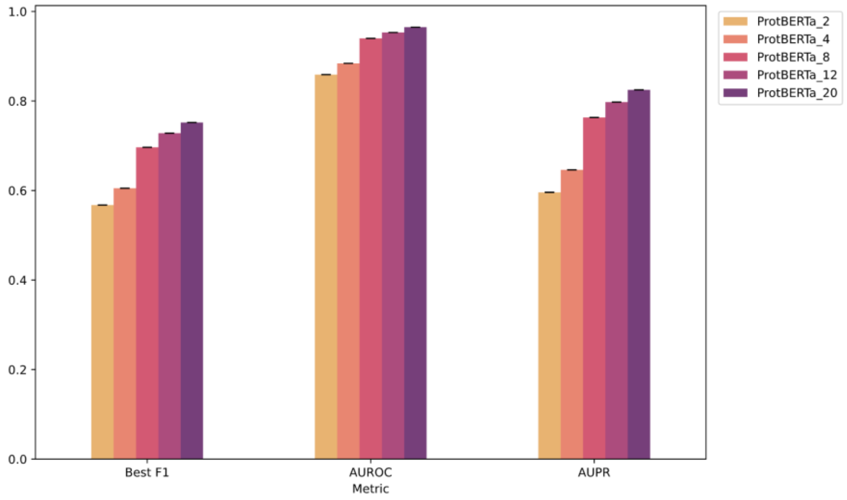


**Supplementary Figure 8. Performance of the five ProtBERTa models on the task of protein-protein interaction prediction.** Standard error is shown as black error bars.

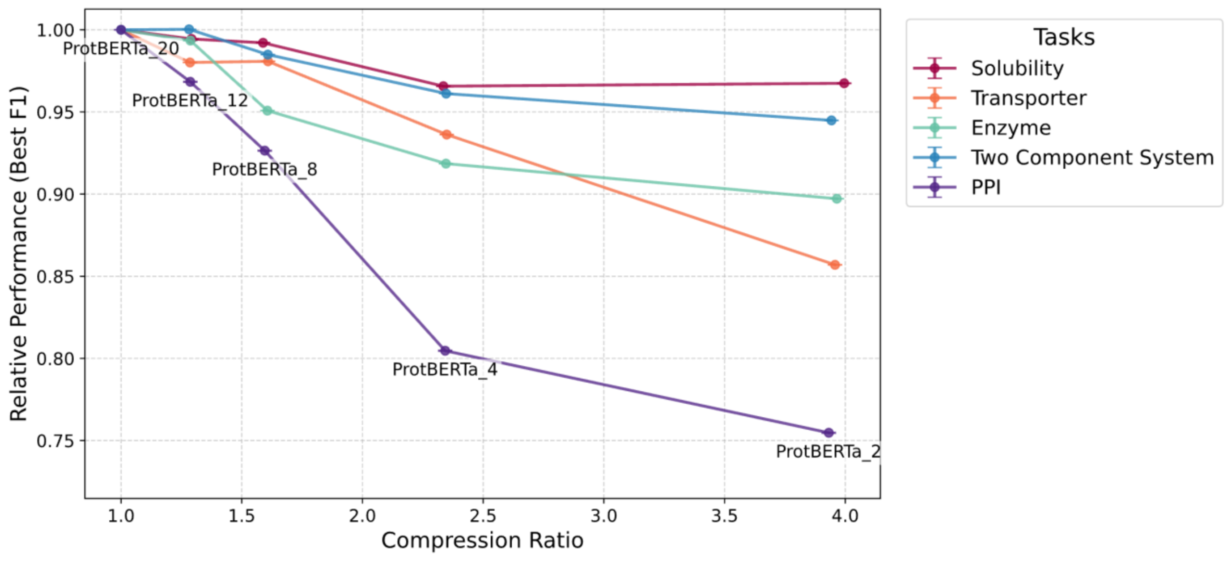


**Supplementary Figure 9. Performance vs. sentence length compression trade-off of the different ProtBERTa models.** The compression is calculated as the average sentence length of each model relative to ProtBERTa_20, and the relative performance is calculated as the best F1 (for the binary tasks) or weighted F1 (for the two-component classification task) relative to ProtBERTa_20.

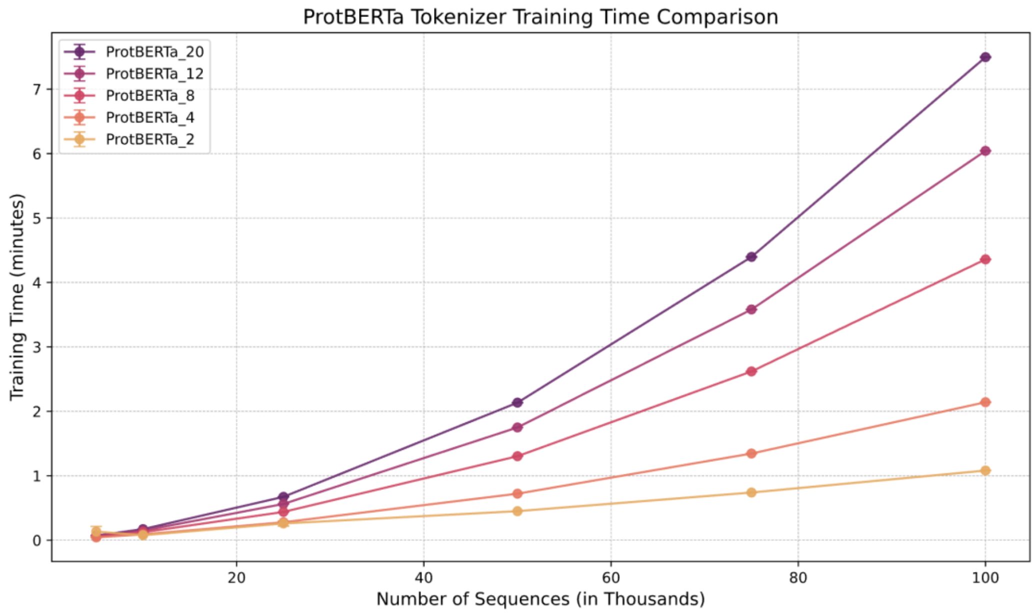


**Supplementary Figure 10. BPE tokenizer training time comparison for the five alphabets.** The training time is measured as a function of the dataset size (in thousands). Error bars represent standard error.

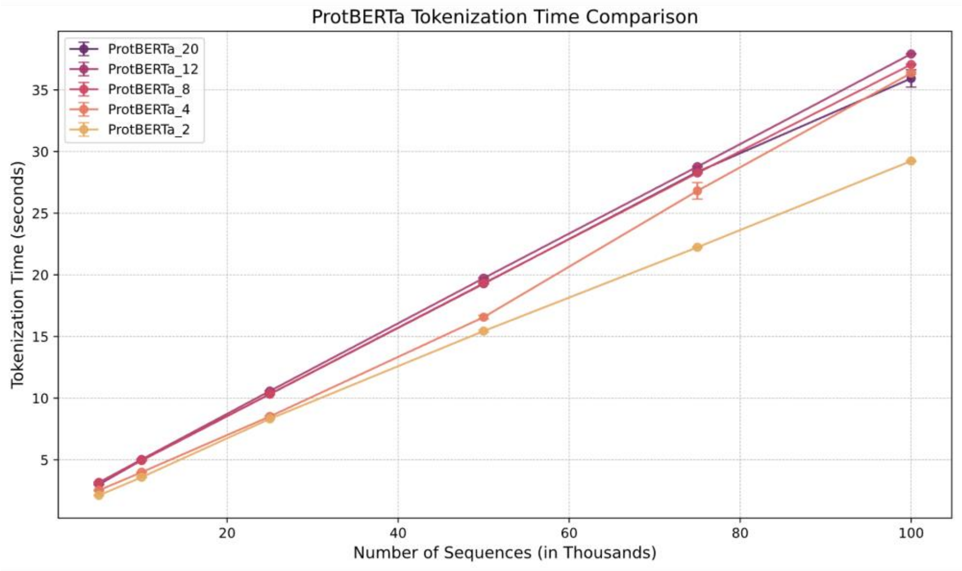


**Supplementary Figure 11. BPE tokenization time comparison for the five alphabets.** The tokenization time is measured as a function of the dataset size (in thousands). Error bars represent standard error.

**Supplementary Table 1. Token and sentence length of the different tokenizers.** Length is presented as mean ± std.

| Model | Token length | Sentence length |
|---|---|---|
| ProtBERTa_2 | 9.89±3.08 | 29.72±25.4 |
| ProtBERTa_4 | 5.74±2.21 | 51.27±44.17 |
| ProtBERTa_8 | 3.88±1.37 | 75.71±65.64 |
| ProtBERTa_12 | 3.1±0.9 | 94.78±81.74 |
| ProtBERTa_20 | 2.4±0.66 | 122.44±106.25 |

**Supplementary Table 2. Description of the amino acid tasks in the DGEB benchmark and their selected performance metrics.**

| Metric Name | Task Description | Metric |
|---|---|---|
| bac_arch_analogs | Identification of functionally analogous protein pairs between Bacteria and Archaea using embeddings cosine similarity. | F1 |
| ModAC_paralogs | Identification of the paralogous ModC protein corresponding to each ModA from a set of orthologous ModC sequences based on cosine similarity. | Recall@50 |
| convergent_enzymes | Enzyme commission (EC) number classification for proteins with no detectable sequence similarity to training proteins from the same class. | F1 |
| EC_numbers | EC number classification with ≤10% sequence identity between proteins of the same class. | F1 |
| biosynthetic_gene_cluster | Multi-label, multi-class classification of proteins according to the metabolites they synthesize. | F1 |
| MopB_clustering | Evaluation of embedding separability of MopB family proteins across catalytic function classes using k-means clustering. | V-measure |
| FeFe_phylogeny | Correlation between pairwise embedding distances and phylogenetic distances derived from tree branch lengths. | Maximum Pearson correlation across cosine, Euclidean, and Manhattan distances |
| RpoB_arch_phylogeny | | |
| RpoB_bac_phylogeny | | |
| Cyano_operonic_pair | Classification of adjacent gene pairs as operonic or non-operonic using cosine similarity of embeddings. | Average precision score calculated using the best threshold |
| Ecoli_operonic_pair | | |
| Vibrio_operonic_pair | | |
| arch_retrieval | Retrieval of functionally analogous bacterial proteins for archaeal or eukaryotic query sequences by ranking bacterial embeddings using cosine similarity. | MAP@5 |
| euk_retrieval | | MAP@5 |

**Supplementary Table 3. Zero-shot Performance of the ProtBERTa models on the homology prediction task**. The best performing model in each metric is marked in bold. Performance is presented with ± standard error on 100 bootstrapped samples.

| Model | AUROC | AUPR | Best F1 |
|---|---|---|---|
| ProtBERTa_20 | **0.896±4e-05** | **0.908±4e-05** | **0.818±6e-05** |
| ProtBERTa_12 | 0.891±5e-05 | 0.902±5e-05 | 0.813±6e-05 |
| ProtBERTa_8 | 0.842±5e-05 | 0.852±6e-05 | 0.767±6e-05 |
| ProtBERTa_4 | 0.764±6e-05 | 0.776±7e-05 | 0.716±6e-05 |
| ProtBERTa_2 | 0.7±6e-05 | 0.72±8e-05 | 0.685±6e-05 |

**Supplementary Table 4. Performance of the ProtBERTa models on the solubility prediction classification task.** The best performing model in each metric is marked in bold. Performance is presented with ± standard error on 1,000 bootstrapped samples.

| Model | AUROC | AUPR | Best F1 |
|---|---|---|---|
| ProtBERTa_20 | 0.921±0.0002 | 0.942±0.0002 | **0.923±0.0002** |
| ProtBERTa_12 | 0.92±0.0002 | 0.94±0.0002 | 0.918±0.0002 |
| ProtBERTa_8 | **0.923±0.0002** | **0.945±0.0002** | 0.916±0.0002 |
| ProtBERTa_4 | 0.913±0.0002 | 0.944±0.0002 | 0.891±0.0002 |
| ProtBERTa_2 | 0.891±0.0003 | 0.915±0.0003 | 0.893±0.0002 |

**Supplementary Table 5. Performance of the ProtBERTa models on the enzyme detection task.** The best performing model in each metric is marked in bold. Performance is presented with ± standard error on 100 bootstrapped samples.

| Model | AUROC | AUPR | Best F1 |
|---|---|---|---|
| ProtBERTa_20 | **0.883±1e-05** | **0.86±2e-05** | **0.804±2e-05** |
| ProtBERTa_12 | 0.877±1e-05 | 0.854±2e-05 | 0.799±2e-05 |
| ProtBERTa_8 | 0.838±2e-05 | 0.811±3e-05 | 0.765±2e-05 |
| ProtBERTa_4 | 0.8±2e-05 | 0.754±3e-05 | 0.739±2e-05 |
| ProtBERTa_2 | 0.767±2e-05 | 0.727±3e-05 | 0.722±2e-05 |

**Supplementary Table 6. Performance of the ProtBERTa models on the transporter detection task.** The best performing model in each metric is marked in bold. Performance is presented with ± standard error on 100 bootstrapped samples.

| Model | AUROC | AUPR | Best F1 |
|---|---|---|---|
| ProtBERTa_20 | **0.954±1e-05** | **0.847±3e-05** | **0.743±4e-05** |
| ProtBERTa_12 | 0.947±1e-05 | 0.823±4e-05 | 0.729±4e-05 |
| ProtBERTa_8 | 0.947±1e-05 | 0.827±4e-05 | 0.729±4e-05 |
| ProtBERTa_4 | 0.936±2e-05 | 0.8±4e-05 | 0.696±4e-05 |
| ProtBERTa_2 | 0.9±2e-05 | 0.697±5e-05 | 0.637±4e-05 |

**Supplementary Table 7. Performance of the ProtBERTa models on the two-component system prediction classification task.** All the metrics are calculated using a macro average across the three classes. The best performing model in each metric is marked in bold. Performance is presented with ± standard error on 100 bootstrapped samples.

| Model | AUROC | AUPR | F1 |
|---|---|---|---|
| ProtBERTa_20 | **0.985±1e-05** | **0.94±4e-05** | 0.901±4e-05 |
| ProtBERTa_12 | 0.983±1e-05 | 0.937±4e-05 | **0.902±4e-05** |
| ProtBERTa_8 | 0.966±2e-05 | 0.885±5e-05 | 0.839±5e-05 |
| ProtBERTa_4 | 0.923±3e-05 | 0.771±7e-05 | 0.74±7e-05 |
| ProtBERTa_2 | 0.886±4e-05 | 0.688±8e-05 | 0.645±8e-05 |

**Supplementary Table 8. Performance of the ProtBERTa models on the protein-protein interaction prediction classification task.** The best performing model in each metric is marked in bold. Performance is presented with ± standard error on 100 bootstrapped samples.

| Model | AUROC | AUPR | Best F1 |
|---|---|---|---|
| ProtBERTa_20 | **0.965±4e-05** | **0.825±0.0002** | **0.752±0.0002** |
| ProtBERTa_12 | 0.953±6e-05 | 0.798±0.0002 | 0.728±0.0002 |
| ProtBERTa_8 | 0.94±7e-05 | 0.763±0.0002 | 0.697±0.0002 |
| ProtBERTa_4 | 0.884±0.0001 | 0.646±0.0002 | 0.605±0.0002 |
| ProtBERTa_2 | 0.859±0.0001 | 0.596±0.0003 | 0.567±0.0002 |

**Supplementary Table 9. Performance of the ProtBERTa models on the stability prediction regression task.** The best performing model in each metric is marked in bold. Performance is presented with ± standard error on 1,000 bootstrapped samples.

| Model \ Metric | ProtBERTa_2 | ProtBERTa_4 | ProtBERTa_8 | ProtBERTa_12 | ProtBERTa_20 |
|---|---|---|---|---|---|
| MSE | 0.5±0.0004 | **0.294±0.0003** | 0.425±0.0004 | 0.556±0.0004 | 0.312±0.0003 |
| MAE | 0.637±0.0004 | **0.474±0.0003** | 0.583±0.0003 | 0.68±0.0003 | 0.481±0.0003 |
| RMSE | 0.707±0.0003 | **0.543±0.0003** | 0.652±0.0003 | 0.745±0.0003 | 0.558±0.0003 |

**Supplementary Table 10. Performance of the ProtBERTa models on the optimal temperature prediction regression task.** The best performing model in each metric is marked in bold. Performance is presented with ± standard error on 1,000 bootstrapped samples.

| Model \ Metric | ProtBERTa_2 | ProtBERTa_4 | ProtBERTa_8 | ProtBERTa_12 | ProtBERTa_20 |
|---|---|---|---|---|---|
| MSE | **308.961±0.419** | 322.61±0.437 | 333.968±0.451 | 369.22±0.487 | 365.052±0.483 |
| MAE | **12.087±0.008** | 12.301±0.008 | 12.451±0.008 | 12.863±0.009 | 12.818±0.009 |
| RMSE | **17.577±0.012** | 17.961±0.012 | 18.275±0.012 | 19.215±0.013 | 19.106±0.013 |

**Supplementary Table 11. Performance of the ProtBERTa models on the fluorescence prediction regression task.** The best performing model in each metric is marked in bold. Performance is presented with ± standard error on 1,000 bootstrapped samples.

| Model \ Metric | ProtBERTa_2 | ProtBERTa_4 | ProtBERTa_8 | ProtBERTa_12 | ProtBERTa_20 |
|---|---|---|---|---|---|
| MSE | 1.823±0.0004 | 1.72±0.0004 | 1.108±0.0003 | **0.86±0.0003** | 1.079±0.0004 |
| MAE | 0.983±0.0002 | 0.947±0.0002 | 0.732±0.0002 | **0.582±0.0001** | 0.688±0.0002 |
| RMSE | 1.35±0.00016 | 1.311±0.0002 | 1.053±0.0002 | **0.927±0.0002** | 1.039±0.0002 |
